# MorphVAE: Generating Neural Morphologies from 3D-Walks using a Variational Autoencoder with Spherical Latent Space

**DOI:** 10.1101/2021.06.14.448271

**Authors:** Sophie Laturnus, Philipp Berens

**Affiliations:** Institute for Ophthalmic Research, University of Tübingen, Germany; Center for Integrative Neuroscience, University of Tübingen, Germany; Tübingen AI Center, Germany

## Abstract

For the past century, the anatomy of a neuron has been considered one of its defining features: The shape of a neuron’s dendrites and axon fundamentally determines what other neurons it can connect to. These neurites have been described using mathematical tools e.g. in the context of cell type classification, but generative models of these structures have only rarely been proposed and are often computationally inefficient. Here we propose MorphVAE, a sequence-to-sequence variational autoencoder with spherical latent space as a generative model for neural morphologies. The model operates on walks within the tree structure of a neuron and can incorporate expert annotations on a subset of the data using semi-supervised learning. We develop our model on artificially generated toy data and evaluate its performance on dendrites of excitatory cells and axons of inhibitory cells of mouse motor cortex (M1) and dendrites of retinal ganglion cells. We show that the learned latent feature space allows for better cell type discrimination than other commonly used features. By sampling new walks from the latent space we can easily construct new morphologies with a specified degree of similarity to their reference neuron, providing an efficient generative model for neural morphologies.

## 1. Introduction

The anatomy of a neuron has fascinated scientists ever since the pioneering work of Cajal (Ramón y Cajal, 1911). The dendritic and axonal processes of a neuron naturally decide what other neurons it can connect to, and thus which inputs it receives and where the computed outputs are sent to (Hill et al., 2012). The anatomical shape of a neuron — its morphology — plays therefore an important role for its function in the circuit. In particular, different types of neurons, and thus different building blocks of the circuit, have fundamentally different morphologies (Markram et al., 2004; DeFelipe et al., 2013).

This variability of neural shapes has been quantified using sets of expert-determined features (Scorcioni et al., 2008; Armañanzas & Ascoli, 2015; Wang et al., 2018; Kanari et al., 2019). While this approach allows classifying neurons into distinct morphological types (m-types), it does not yield a generative model for new neurons in a straightforward manner. Algorithms for generating neurons have been suggested which either start with simple neuron shapes and assume a distinct set of biologically motivated growth rules (van Pelt & Schierwagen, 2004; Eberhard et al., 2006; Bingham et al., 2020; Kassraian-Fard et al., 2020) or manipulate these shapes to iteratively match a set of properties from observed data (Cuntz et al., 2011; Serene, 2013; Farhoodi & Kording, 2018).

However, a unified, efficient framework for modeling the cell type diversity of large sets of neuron morphologies and generating new morphologies has been missing. Here we propose MorphVAE, a sequence-to-sequence (seq2seq) variational autoencoder with spherical latent space (Davidson et al., 2018; Xu & Durrett, 2018) working on 3D-walks along a neuron’s morphology. The generated morphologies match key characteristics of their biological counterparts even though no biological constraints are incorporated in the model. Furthermore, MoRPHVAE yields a feature representation which is at least as good or better than state-of-the-art morphology representations.

## 2. Methods

Our goal is (1) to build a generative model of a diverse set of realistic looking neural morphologies and at the same time (2) to learn a latent representation of neuron morphologies revealing cell type related differences. Our model is trained on a set of neural morphologies represented by their tree graph *T*, possibly with assigned cell type label *c_T_*. For each individual reference neuron, the model operates on 3D-walks along the neurites (Fig. 1).

**Figure 1.**
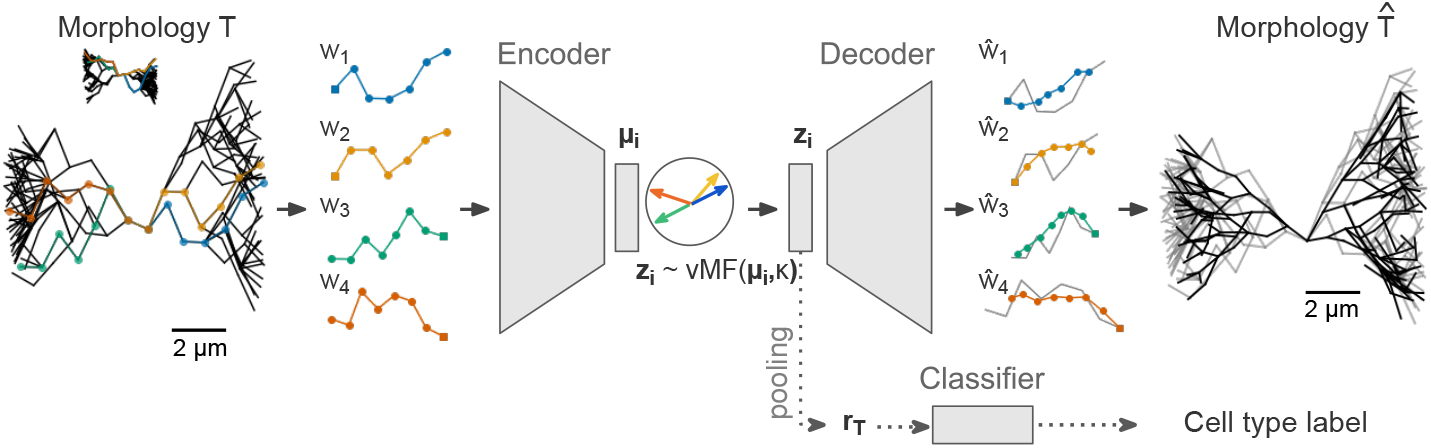
From a given morphological tree graph *T* (*left*) we sample walks *w_i_* from the soma to the tips. These walks are used to train a variational seq2seq-autoencoder with a multivariate von-Mises Fisher distributed latent space with fixed variance *κ*. Each walk encodes a mean direction *μ_i_* in the latent space that is used to sample a latent variable *z_i_* ~ vMF(*μ_i_, κ)* (*middle*). A subsequent decoder decodes *z_i_* trying to match the input walk. The decoded walks 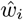 are then clustered to construct a new tree graph 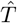 (*right*). The black lines show the reconstruction while the reference neuron is shown in grey. A logistic regression classifier that pools over the latent samples *z_i_* of all walks *w_i_* within one neuron can be used to inform the model about labels.

In the following section, we first describe how the 3D-walks are obtained (Section 2.1) and why they are advantageous to using a neuron’s tree graph directly. Second, we present a generative model (Section 2.2) that will generate a set of new 3D-walks for a given reference neuron. Additionally, we explain how we use the encoded walks in the latent space to obtain a feature representation for a neuron that can incorporate expert annotations in a semi-supervised fashion (Section 2.3). Finally, in Section 2.4 we describe how we construct a new tree graph 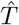 from the generated set of walks with minimal biological constraints.

### 2.1. Sampling 3D-walks along neurites

A neural morphology is represented as a directed tree graph defined as a tuple *T* = (*V, E*). The first entry *V* is the set of nodes, 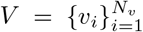, where 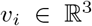 represent coordinates in 3D space and *υ*_1_ = (0,0,0) denotes the neuron’s soma which is the root of the tree. The second entry *E* = {*e_ij_* = (*υ_i_, υ_j_*)|*υ_i_, υ_j_* ∈ *V*} is the set of directed edges which connects two nodes in *V*. The leaf nodes which have no outgoing edge are called tips and denote as 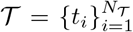. We define *w* = (*υ*_1_, *υ*_*i*_1__, *υ*_*i*_2__, …, *t_k_*) as a finite 3D-walk from soma *υ*_1_ to tip *t_k_* where each pair of consecutive nodes (*υ_i_j__, υ_i_l__*) is connected via an edge *e_i_j___i_l__*. Given a morphology graph *T*, we can represent it as the set 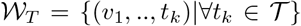 which contains all existing walks from soma to tip.

A key advantage of this representation is that it describes a neuron’s geometry and its topology at the same time in such a way that allows us to leverage seq2seq models. The succession of two coordinates within one walk implies a connecting edge in that direction. Because all walks start at the soma and progress outwards to the tips the underlying morphology can be reconstructed given enough walks have been sampled.

Note that the number of nodes and the number of walks and their length varies from neuron to neuron. To speed up model fitting, we fix the length *l* of each walk and randomly sample *n_w_* = 256 walks with replacement from each 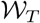 to obtain a matrix 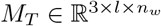; longer walks are truncated and shorter walks are padded with zeros and packed using the `pack_padded_sequence` utility of PyTorch (Paszke et al., 2019). We use the set of *N* pairs 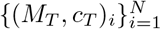 consisting of a walk representation and a cell type label, to train the MorphVAE model.

### 2.2. Generative model

To model the distribution over walks from soma to tip within a neuron, we employ a seq2seq variational autoencoder with spherical latent space (Sutskever et al., 2014; Xu & Durrett, 2018; Davidson et al., 2018), a model that has been originally developed in the context of natural language processing which we modify here to predict continuous variables. Our goal is to find an encoder *f_θ_*(*z*|*w*) for the walk *w* with 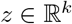, 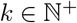, and a decoder *g_ϕ_*(*w*|*z*) such that *g_ϕ_*(*f_θ_*(*w*)) ≈ *w*.

Here, *f_θ_*(*z*|*w*) = vMF(*μ, κ* = *c*) is a von-Mises Fisher (vMF) distribution with fixed variance *κ* whose mean *μ* is modelled by a two-layered unidirectional Long Short-Term Memory (LSTM) unit (Hochreiter & Schmidhuber, 1997) with linear input and output layer. A LSTM is a recurrent neural network that can keep track of already seen input via two internal states, its hidden state *h*, and its cell state *c*.

In the encoder *f_θ_*(*z*|*w* = (*υ*_1_, …, *υ_n_*)), each coordinate *υ_i_* is first projected into a higher dimensional space via a linear transformation *x_i_* = *W_in_* · *υ_i_* with 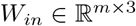. The *x_i_* are then consecutively fed through a unidirectional two-layered LSTM, whose internal states have been initialized to 0^*m*^, until we obtain (*h_n_, c_n_*) = LSTM(*x_n_, h_n-1_, c_n-1_*). The last hidden and cell state of both layers is then concatenated (denoted by 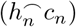) and linearly projected onto *k* dimensions using 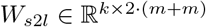, the transformation matrix converting from LSTM states to the latent space, to obtain the mean of our vMF distribution for the sampling of *z*. Finally, *z* is taken as the average over five samples of *z_i_* ~ vMF(*μ, c*) which are sampled via rejection sampling (Xu & Durrett, 2018; Davidson et al., 2018).

In summary:

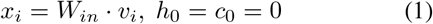

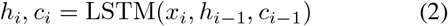

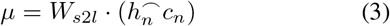

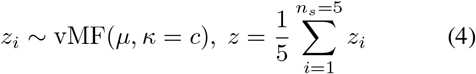

The decoder *g_ϕ_*(*w*|*z*) decodes the coordinates 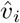 in 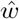 step by step from *z* and 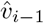 using a second unidirectional two-layered LSTM. First, the LSTM is initialized with the initial states 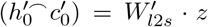, where 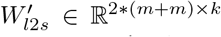 transforms *z* back into LSTM state space, and a linear projection of the first coordinate 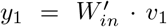 with 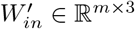. Then, the LSTM subsequently predicts *y*_*i*+1_ from its internal states and the previous *y_i_*. Finally, each *y_i_* is passed through a linear transformation 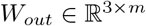 to predict the output sequence 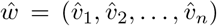. In summary:

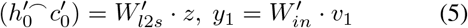

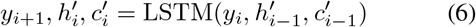

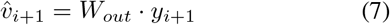

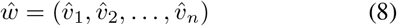

We jointly optimize the parameters {*θ, ϕ*} of the en- and decoder by maximizing the evidence lower bound (ELBO)

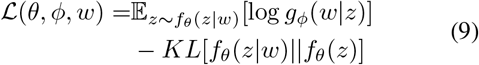

using a uniform vMF prior *f_θ_*(*z*) = vMF(·, 0) on the latent space. The vMF prior prevents the *KL collapse* typically observed in Gaussian VAE settings (Bowman et al., 2015). In fact, the KL term in our loss term is constant and only depends on the chosen variance *κ* (Xu & Durrett, 2018). Thus, we can simplify Eq. 9 to

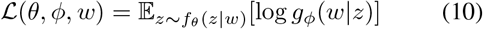

which we estimate by the summed mean-squared error between *w* and 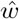.

### 2.3. Learning a semi-supervised neuron representation

From the learned latent space over 3D-walks we additionally derive a feature representation *r_T_* for each neuron *T*. For this, we pool over the *n_w_* = 256 walks in *M_T_*, now encoded and sampled from the latent space 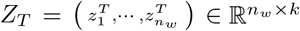. Thus,

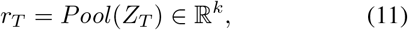

where *Pool* denotes a pooling operation over all walks, like max-pooling or averaging, that is insensitive to the order of the walks in *Z_T_*. We explored different pooling operations during training (for details, see Appendix).

To incorporate information from potentially available cell type labels, we added a classification head that predicts the cell type label 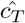 of *r_T_* using logistic regression (Fig. 1). The classifier is tied to the autoencoder via the latent sample 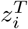 of each walk in *T* and is jointly trained to minimize the unweighted cross-entropy loss. This allows us to incorporate knowledge about cell type labels when learning the representations for the walks, *μ_i_*, and for the neurons, *r_T_*. Notably, this approach allows to integrate any label information that is relevant to the researcher. For our experiments, we vary the fraction of cells with labels provided to train the model, interpolating between an unsupervised, a semi-supervised and a fully supervised setting. All parts of the model were implemented in PyTorch (Paszke et al., 2019), all code is available at https://github.com/berenslab/morphvae.

### 2.4. Sampling of morphologies

For a given reference morphology *T* we want to construct a new morphology 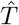 that resembles *T* with respect to type-defining morphological properties. To this end, we pass the walks *w_i_* in the matrix *M_T_* through our model as described in Section 2.2 to obtain a matrix

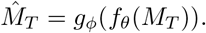

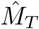 is a noisy version of *M_T_*, however, from which we cannot reconstruct a new morphology directly.

First, we need to estimate the proper walk length of walks that are shorter than *l*. We cannot use an end-of-sentence token due to the continuous nature of our setup, and we employed zero-padding, thus shorter walks jump back to (0,0,0). Here, we trimmed each walk *w* whenever its path angle, the angle between two consecutive segments, exceeded 75 degree.

Second, we reduced the number of nodes by aggregating nodes before constructing a new neuron tree as, otherwise, we would over-estimate the number of nodes 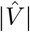 in 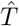 and thus overestimate the number of dendrites. To this end, we clustered each column 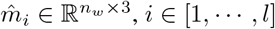, in 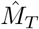 separately from last to first step using a fixed distance threshold *d* (*d_toy_* = .5, *d_m1exc_* = .4, *d_m1inh_* = .3, *d_rgc_* = .25). For clustering, we used agglomerative clustering as implemented in `scikit-learn` (Pedregosa et al., 2011) with a Ward linkage criterion (Ward Jr, 1963) and a Euclidean distance metric. In some cases, this will result in walks that have been merged in step *i* to be split again in step *i* – 1. If this happens, we merge the involved clusters at step *i* – 1 to avoid illegal paths. Now, we replace each coordinate in 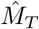 with its respective cluster mean to obtain a clustered matrix 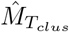.

Finally, we construct a new tree 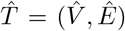 by ‘reverse engineering’ the walks in 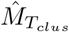:

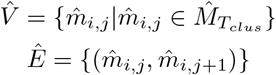

for *i* ∈ [1, …, *n_w_*] and *j* ∈ [1, …, *l*].

### 2.5. Datasets

We trained and evaluated the MorphVAE model on four different datasets: artificially generated toy data, excitatory pyramidal cell dendrites in M1 (Scala et al., 2020), inhibitory cell axons in M1 (Scala et al., 2020), and retinal ganglion cell dendrites (Reinhard et al., 2019), where all data was recorded from adult mice (Fig. 2).

**Figure 2.**
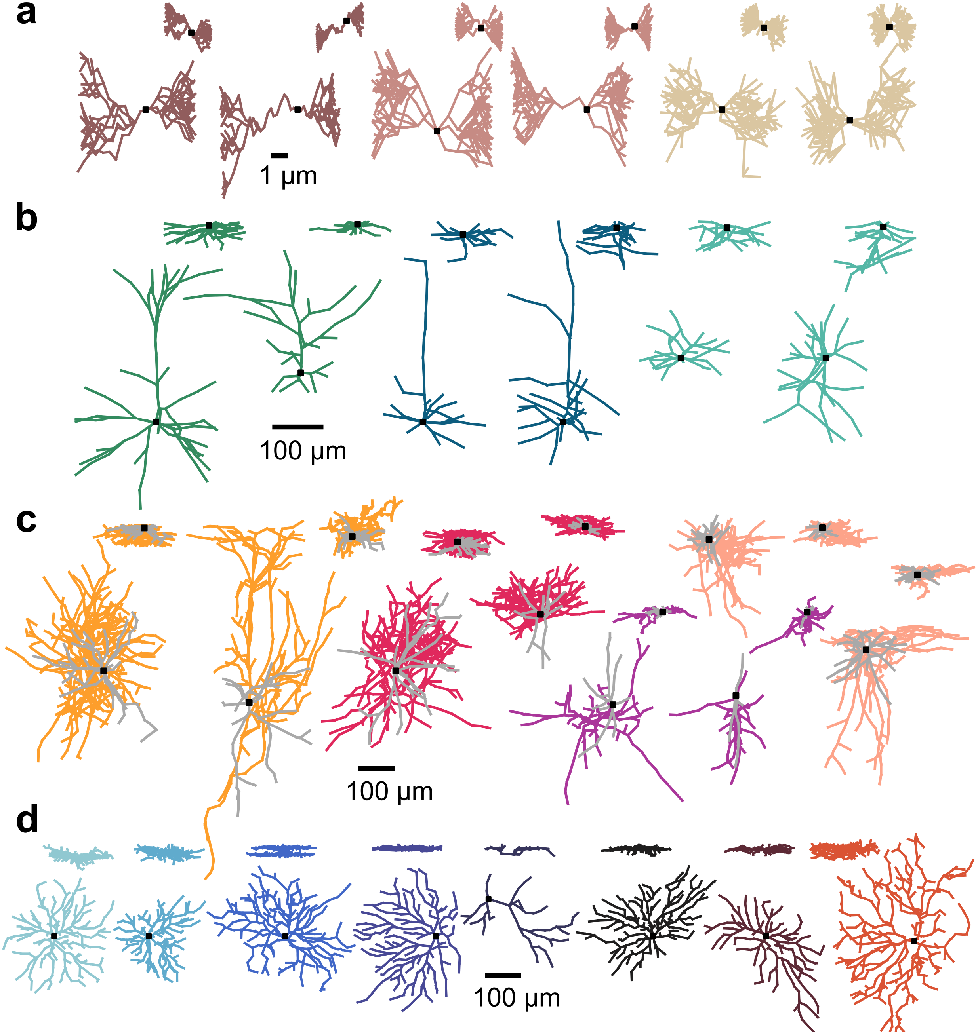
A random subset of two morphologies for each population within each dataset. **a**) Artificially generated data (*brown*: P1, *antique pink*: P2, *beige*: P3, inset: *xz* view). **b**) Excitatory cell dendrites from mouse M1 (green: Tufted, *dark blue*: Untufted, *light blue*: Others, inset: *xy* view). **c**) Inhibitory cells from mouse M1. Axons are in color, dendrites are in grey (*yellow*: Sst, *pink*: Pvalb, *purple*: Vip, *rose*: Lamp5, inset: xy view). **d**) One example for eight out of 14 populations of retinal ganglion cell dendrites (inset: xz view). Somata are indicated by a black square.

#### 2.5.1. Artificial dataset

We generated a set of *N* = 1200 artificial neurons using very simple growth and branching rules (for details, see Appendix). The resulting toy neurons were not “real” in the sense that they mimicked actual neurons accurately, but they did resemble the overall shape and branching patterns of real neural populations. Each neuron contained |*V*| = 200 nodes and belonged to one of three different populations *P_i_* of equal size (*N_P_i__* = 400) where each population had its unique set of generating parameters (Fig. 2**a**). These were chosen such that the neuron populations were not trivial to separate e.g. by PCA on density maps. For the MorphVAE model, we set the walk length to *l* = 16 when generating the walk matrices *M_T_i__* and split the data into *N_train_* = 750, *N_val_* = 250, and *N_test_* = 200.

#### 2.5.2. Real neuron datasets

We downloaded 275 dendritic reconstructions of excitatory neurons and 372 axonal reconstructions of inhibitory neurons^1^ that had been recorded in a large scale multi-modal study describing cell types in adult mouse M1 (Scala et al., 2020). We manually assigned the m-type of the excitatory neurons to one of tufted, untufted or other based on visual inspection of the apical dendrites (for details, see Appendix). For the inhibitory neurons we used the assigned RNA family labels (t-type) as cell type labels but grouped Sncg to Vip as it contained only 6 cells. We also downloaded 599 reconstructions of retinal ganglion cell dendrites from neuromorpho (Ascoli et al., 2007) that were originally collected by Reinhard et al. (2019). Here, we used the cell type labels assigned by the authors which were based on the cells’ stratification pattern within the inner plexiform layer.

All real reconstructions were soma centered and resampled to reduce the number of nodes within each reconstruction (sampling distance see Table 1). The 3D coordinates were rescaled by a factor of 100 to allow transfer learning from the MorphVAE trained on artificial data. We set the walk length to *l* = 32 when generating the walk matrices *M_T_i__* and split the data into a stratified training, validation, and test set of the sizes reported in Table 1. Due to the imbalance in class sizes we report balanced accuracy 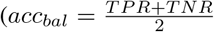, with 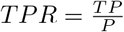 and 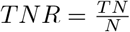 for the analyses in Section 3.2. For a more detailed account of each data set, see the Appendix.

**Table 1.** Number of classes *N_P_*, resampling distance, and sizes of stratified training, validation, and test sets for each dataset.

| Dataset | $N_P$ | resampled at | $N_{train}$ | $N_{val}$ | $N_{test}$ |
| --- | --- | --- | --- | --- | --- |
| Toy | 3 | – | 750 | 250 | 200 |
| M1 EXC | 3 | 50 μm | 160 | 55 | 60 |
| M1 INH | 4 | 40 μm | 248 | 62 | 62 |
| RGC | 14 | 30 μm | 400 | 99 | 100 |

### 2.6. Training

On the artificial dataset we fit the entire model over 150 epochs using the Adam optimizer (Kingma & Ba, 2014) with a batch size of 128 and an initial learning rate of 0.01 that we half at every 50 epochs. Additionally, we employed teacher forcing (Lamb et al., 2016) at a rate of 50% to train the decoder and we regularized the model using dropout (Srivastava et al., 2014) in the input layers of the encoder, the decoder and the classification head, and in both LSTMs. To find the optimal values for the network dimensions and pooling operations we performed a grid search and took the model with the best average validation performance over three Glorot initializations (Glorot & Bengio, 2010) (for details, see Appendix). The best performing model used a hidden and a latent dimension of *m* = *k* = 32, a variance of *κ* = 500 and max-pooling. On the M1 data, fitting the model from scratch was unsuccessful, probably due to the low sample size. Thus, we first fit the model as described above but on an artificial dataset that included more diverse neurons (for details, see Appendix) and employed random scaling as that improved the quality of the reconstructions. We then fine-tuned this model for another 200 epochs (without random scaling) on each of the M1 datasets. The model sucessfully trained on the RGC data without pre-training, but pre-training led to better results. We therefore report the results of the pre-trained models throughout.

### 2.7. Embeddings, classification and density maps

To visualize the encoded representation of walks, *μ_i_*, in two dimensions we used *openTSNE* (Poličar et al., 2019), an open and fast implementation of t-distributed stochastic neighbor embedding (t-SNE) (Van der Maaten & Hinton, 2008), with PCA-initialization, cosine distance and a perplexity of 200 (Kobak & Berens, 2019). For the embedding of the neuron representations *r_T_i__* we used a perplexity of 30. We show the embeddings on test data for the model with the best performance in each condition unless stated otherwise.

To evaluate the neuron representation using a ‘upper bound on discriminability’, we used a *k*-nearest neighbor classifier (*k* = 5) as implemented in `scikit-learn` (Pedregosa et al., 2011). We fit the classifier on the representations *r_T_i__* of the training data for each of the three model initialization and evaluated on the respective encoding of the test data. We report averages across the three initializations throughout.

We computed density maps and morphometric statistics using the `MorphoPy` toolbox (Laturnus et al., 2020b). For the density maps, we sampled equidistant points with 0.1 μm (toy data) and 1 μm (all other data) spacing along each neurite of *T* and normalized the resulting point cloud to lie between 0 and 1. We chose the normalization ranges globally within each dataset to preserve relative sizes between cells. The normalized point cloud was then projected onto the cardinal planes or axes (toy data: *xy*, M1 EXC/M1 INH: *xz*, RGC: *z*), and binned into 20 equidistant bins along each direction. We smoothed the resulting histograms by convolving them with a 11-bin Gaussian kernel with a varying standard deviation of *σ* ∈ [.5,1, 2] bins to find the best projection. We treated the density maps as flattened vectors and reduced them to as many principal components (PC) as needed to keep more than 95% of the variance (for details, see Appendix). We used the morphometric statistics as they are implemented in the toolbox per default.

### 2.8. TREES Toolbox

To compare our generative model with existing work, we generated reconstructions of all test set neurons in each real dataset using the TREES Toolbox (Cuntz et al., 2011). For this we generated a 3D image stack of each neuron (for details, see Appendix), passed them through a custom MATLAB script, and sampled one new morphology per stack.

## 3. Results

### 3.1. Model performance on the artificial dataset

#### 3.1.1. Training and ablation study

First, we validated the model on the artificial dataset. We started with a fully supervised setting, where the model attempted to reconstruct the neural morphologies and classify the neurons correctly based on their latent representation *r_T_*. In this setting, the classification head was fully trained after 100 epochs and its loss plateaued while the autoencoder still improved its performance. One run over 150 epochs on the 750 training morphologies took about 2 hours on one NVIDIA Titan Xp GPU with 12 GB memory.

To investigate if the model can also be trained in a semi- or unsupervised setting, we systematically changed the amount of labels provided to the classification head during training from 100% to 0% of labels, moving from a fully supervised to a semi-supervised and unsupervised setting (Table. 2). We noticed that the reconstruction loss changed from 519.4 ± 14.5 to 575 ± 11.7 (mean ± standard deviation across three initializations) when not allowing access to the cell type labels, indicating that label information also helped the generative model to create better walk reconstructions.

We studied the influence of training set size on model performance by reducing the amount of training data in steps of 150 samples. Hereby, the model reconstructed reasonably well until *n* = 450 (617.1 ± 11.7; mean ± SD) and then deteriorated quickly (Table 2).

**Table 2.** Reconstruction and classification loss on the test set when ablating different parameters during training on the artificial dataset. Values denote the mean ± standard deviation across three model initializations.

| Model | Rec-Loss | Class-Loss |
| --- | --- | --- |
| Standard | **$519.4 \pm 14.5$** | **$19.0 \pm 1.3$** |
| n=600 | $586.3 \pm 3.8$ | $24.4 \pm 1.$ |
| n=450 | $617.1 \pm 11.7$ | $30.8 \pm 4.5$ |
| n=300 | $793.5 \pm 61.1$ | $34.4 \pm 1.$ |
| n=150 | $1268.4 \pm 47.3$ | $64.9 \pm 6.1$ |
| Shuffled | $557.5 \pm 4.$ | $291.4 \pm 2.$ |
| No labels | $575.4 \pm 11.7$ | $71.6 \pm 2.6$ |
| l=8 | $3737.8 \pm 295.6$ | $37.1 \pm 11.$ |
| l=32 | $583.4 \pm 6.1$ | $25.7 \pm 2.8$ |

We also varied the walk length *l* to assess its influence on the model performance. If *l* is chosen too short, the structure of each neuron cannot be accurately sampled, if it is chosen too long it creates strong zero-padding in the walk matrix *M_T_*. Both settings were harmful to the model but choosing it too short was more severe (Table 2).

#### 3.1.2. Learned neuron representations in MorphVAE

We next studied the quality of the learned neural representation *r_T_*. To this end, we created a 2D visualization of all *r_T_i__* in the test set using t-SNE (Van der Maaten & Hinton, 2008). This revealed three distinct, well separated clusters for each population, especially if a high fraction of labels was used (Fig. 3**a**). The separability slowly decreased with decreasing number of cell type labels until the cluster for population *P*2 and *P*3 started to merge (Fig. 3**a**). Yet, even a moderate amount of labels yielded very accurate representations of the three cell types, and also without labels, the representations of the three types did not overlap.

**Figure 3.**
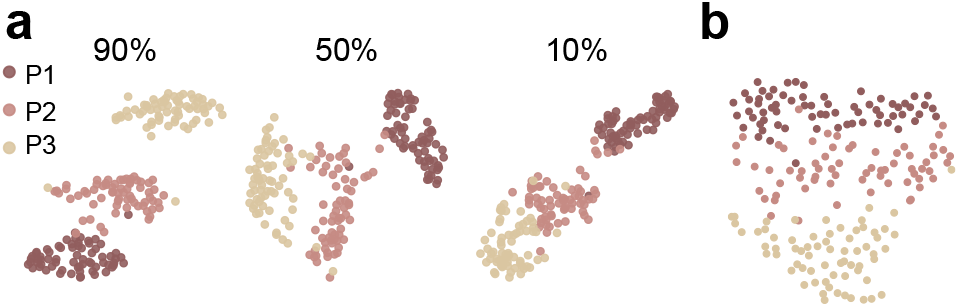
T-SNE embeddings of the neural representations *r_Ti_* for the artificial data for the best performing models using 90%, 50% or 10% of labels during training. **b**) Same as in **a** using the first 10 PCs of *XY* density maps as neural representation.

Additionally, we quantified the discriminability of the learned neural representations training a 5-nearest neighbor classifier on the respective features when using different amounts of the labels (100%, 90%, 50%, 10%, 0%). The prediction accuracy on test data was close to perfect when all labels were used (98% ± 0%, mean ± SEM across initializations) and worsened only slightly for the fully unsupervised case (94% ± 2%).

We compared our findings to the performance when using *XY* density maps as a predictor of cell type label. Density maps project the neural point cloud onto a plane or an axis and have been a classical descriptor in cell typing studies (Jefferis et al., 2007; Sümbül et al., 2014; Laturnus et al., 2020a). We found that the MorphVAE representation had much better separability both in terms of visualization (Fig. 3**b**) and in terms of prediction accuracy (Table 3).

**Table 3.** Balanced classification accuracy on the test set using the learned neuron representation *r_T_* (mean ± SEM across initializations) and the best competing density map. The values in brackets indicate the amount of labels used during training or the width of the smoothing kernel for density map generation.

| Dataset | $N_P$ | Representation | Accuracy |
| --- | --- | --- | --- |
| Toy | 3 | $r_T$ (100%) | **98% ± 0** |
| | | $r_T$ (0%) | 94% ± 2 |
| | | $DM_{xy}(\sigma = 2)$ | 90% |
| M1 EXC | 3 | $r_T$ (100%) | **70% ± 5** |
| | | $r_T$ (0%) | 58% ± 7 |
| | | $DM_{xz}(\sigma = 1)$ | 60% |
| M1 INH | 4 | $r_T$ (50%) | 56% ± 8 |
| | | $r_T$ (0%) | 52% ± 7 |
| | | $DM_{xz}(\sigma = 1)$ | **66%** |
| RGC | 14 | $r_T$ (90%) | 51% ± 6 |
| | | $r_T$ (0%) | 33% ± 5 |
| | | $DM_z(\sigma = 1)$ | **53%** |

#### 3.1.3. Walk latent space encodes walk length and direction

We also explored the structure of the learned latent space for 3D-walks using 2D visualizations with t-SNE. This space was highly ordered and contained information about the walk length and general direction of each walk in terms of *x*-, *y*-, and *z*-coordinates of its tip (Fig. 4**a–d**). Also, our cell type labels changed gradually over the walk representation (Fig. 4**e**) which explains the emergence of the separated neuron representation even in the unsupervised case.

**Figure 4.**
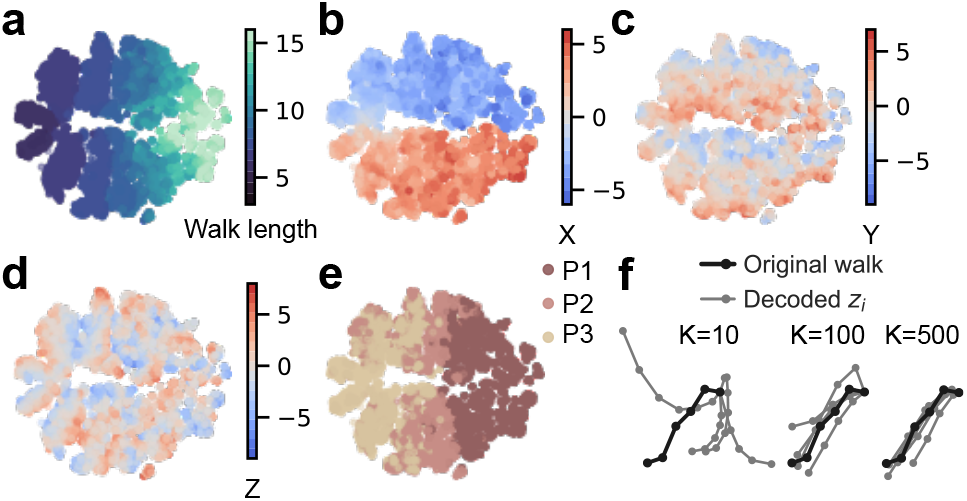
T-SNE representation of the encoded walks *μ_i_* for the artificial data colored by walk length (**a**), by *x*-value (**b**), by *y*-value (**c**) and by *z*-value (**d**) of the tip as well as by cell type label (**e**) of each neuron that the walk was sampled from. **f**) Original walk (black) and five decoded samples (grey) using different variances during the sampling in the vMF latent space. A low *κ* induces a high variance and vice versa.

The reconstruction performance for single 3D-walks was good, with reconstructed walks being slightly smoothed as to be expected from MSE loss (Fig. 4**f**). Additionally, we were able to control the faithfulness of the reconstruction by varying the variance *κ* during sampling in the vMF latent space. For large *κ*, i.e. small variance, the reconstructions were close to the input while small *κ* resulted in larger deviations (Fig. 4**f**).

#### 3.1.4. Sampling morphologies with MorphVAE

Finally, we sampled new morphologies from reference neurons in our dataset using MorphVAE (Fig. 5**a**). For this, we encoded the walk matrix of each reference neuron in the latent space and sampled new walk matrices as described in Section 2.4 using three different sampling variances (*κ* ∈ [100,300,500]). The resulting morphologies agreed well in overall shape and closely matched the observed distributions for certain morphometric statistics (e.g. maximal branch orders, mean soma exit angle and tree asymmetry; Fig. 5**b**, upper row). However, geometric features like the width, and the depth or branching angles within the morphological tree were consistently underestimated, especially in population *P*3 (Fig. 5**b**, lower row), which might be related to the smoothing properties of MSE. In this dataset there was no *κ* that was clearly superior to the others in terms of matching the observed morphometrics but, as expected, higher *κ* yielded narrower distributions (Fig. 5**b**).

**Figure 5.**
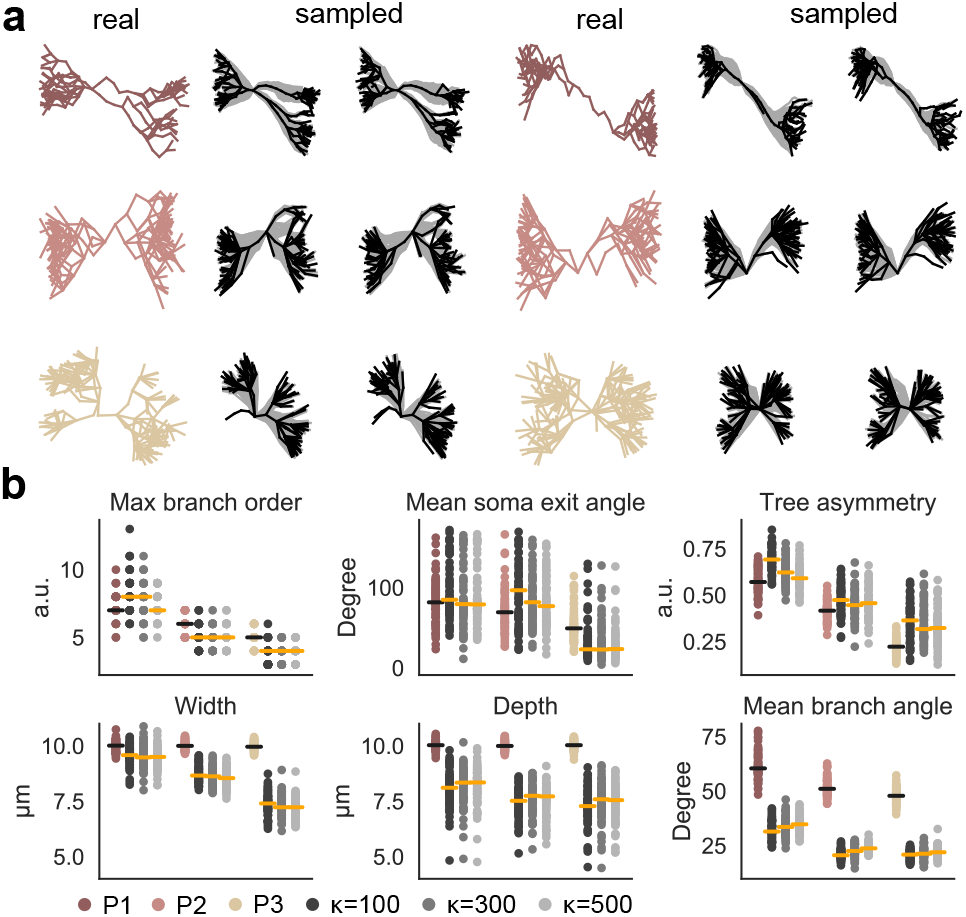
**a**) Ground truth examples and two new samples thereof (*κ* = 500) for each of the three populations. The underlying unclustered 3D-walks are shown in grey. **b**) Distributions of selected morphometric statistics for the test neurons in each population (colored) and the sampled neurons using different values of *κ* during sampling in the latent space (grey). Lines indicate the medians.

### 3.2. MorphVAE on real data

After we validated our approach on the artificial data, we applied the MorphVAE model on three diverse real datasets of neural reconstructions: excitatory pyramidal cell dendrites (M1 EXC, Fig. 2**b**), inhibitory cell axons (M1 INH, Fig. 2**c**), and retinal ganglion cell dendrites (RGC, Fig. 2**d**). We employed transfer learning where the model was first trained on an adjusted artificial dataset with random scaling and then fine-tuned on the real data. This was necessary as both M1 datasets were too small to be trained from scratch but this also improved the reconstruction accuracy for the RGC data (48.9 ± 4.13 vs 67.6 ± 1.77).

In contrast to the artificial data, the reconstruction loss was best when using no labels during training (M1 EXC: 66.42± 2.28, mean ± SD across initializations; M1 INH: 203.94 ± 5.22; RGC: 48.9 ± 4.13) but the fully supervised setting was only slightly worse (M1 EXC: 83.97 ± 10.34; M1 INH: 235.83 ± 11.74; RGC: 74.56 ± 3.05). Nevertheless, this might indicate that classification and reconstruction denote competing tasks on the latent space for real data.

We used a 5-nearest neighbor classifier to predict the cell type labels on the neural representations learned with different amounts of labelled data and reported the balanced classification accuracy on the test data for each dataset (Table 3). The accuracy was good for the representations of excitatory neurons (70% ± 5; mean ± SEM) and better than the best performing density map (60%). For the inhibitory neurons and the RGCs, the accuracy was clearly above chance level but here density maps performed slightly better. Interestingly, for the inhibitory neurons, the best performing representation used only 50% of labelled data (56% ± 8) and was just marginally better than the fully unsupervised representation (52% ± 7). Nevertheless, using label information during training improved the structure in the neuron latent space generally (Fig. 6**a, c**, and **e**), and for RGCs in particular where a clear separation emerged between ON and OFF cells (Fig. 6**e**).

**Figure 6.**
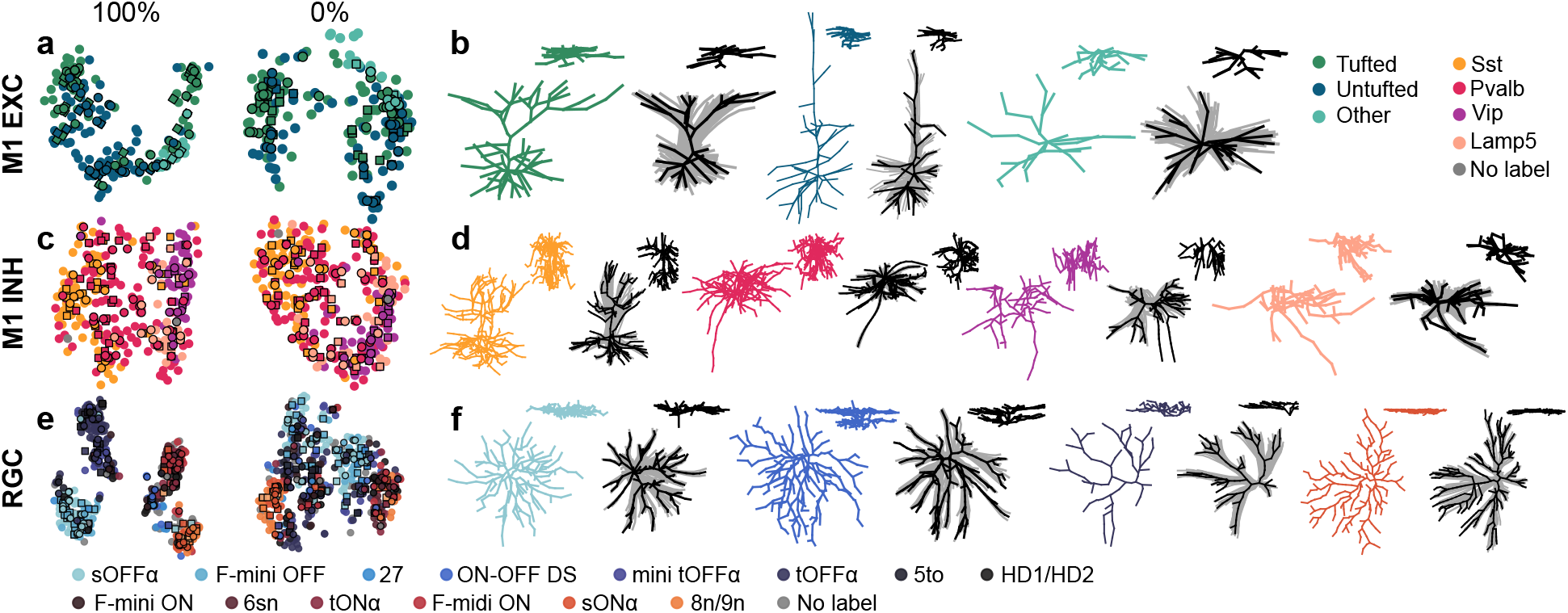
**a**) T-SNE embeddings of the neural representations *r_T_i__* for the dendrites of excitatory cells in M1 for the best performing models using 100% and 0% of labels during training. We create the embedding on the training data (no edges) and project the validation data (squares, black edges) and the test data into it (black edges; *k* = 5, perplexity=5). **b**) One ground truth example and one new sample thereof (*κ* = 500, inset: *xy*) for each of the three excitatory populations. The underlying un-clustered 3D-walks are shown in grey. **c**) T-SNE embedding for axons of inhibitory cells in M1 as in **a**. **d**) One ground truth example and one new sample thereof (*κ* = 500, inset: *xy*) for each of the four inhibitory populations. **e**) T-SNE embedding as in **a**) for the retinal ganglion cell dendrites. **f**) One ground truth example and one new sample thereof (*κ* = 500, inset: *xz*) for four example neurons.

Newly sampled morphologies matched their reference neurons well in overall shape for all three datasets, and often regardless of the cell type label (Fig. 6b, d, f). The model successfully learned to recreate tufted and untufted excitatory cells, for example, or the prominent bistratification of ON-OFF direction selective RGCs (Fig. 6 **f**). Investigating the morphometric statistics of the newly sampled data revealed that pre-training with random scaling ameliorated the underestimation of neural size as width, depth, and height were now comparable with those of the true populations (data shown in Appendix). Hereby, *κ* > 100 generally created better results, yet, the model consistently underestimated the mean branch angles and the median path angles, and created nodes with an unrealistic high degree of branching which we believe to be related to the post-hoc clustering of the sampled walk matrices.

We also created new samples of the same neurons using the TREES Toolbox (Cuntz et al., 2011) (Section 2.8) and extracted their morphometric statistics for comparison with MorphVAE (data shown in Appendix). While the overall match was good for TREES, it often generated narrower distributions for each statistic than what was indicated by the ground truth data.

Finally, we compared the runtime during morphology sampling for the MorphVAE model and the TREES Toolbox in MATLAB Online. Hereby we constrained our system to the same amount of CPUs as provided by MATLAB (16 CPUs) and disabled GPU processing for a fair comparison (for details, see Appendix). The runtime was similar for both models during sampling and tree construction (Ours: 0.53s ± 0.03, TREES: 0.64s ± 0.64; mean ± SD), but differed considerably during data loading (Ours: 6.77s ± 0.83, TREES: 16.49s ± 1.19). Note, that the TREES Toolbox needs to load a new image stack for each neuron separately while MorphVAE encodes the reference walk matrices once for each batch which gives it a strong computational advantage. MorphVAE additionally allows GPU processing which cut the encoding time in half (Ours: 2.94s±0.01).

Thus, MorphVAE denotes an efficient generative model that can be used to sample realistic looking neurons from a diverse set of examples while yielding informative lowdimensional embeddings at the same time.

## 4. Discussion

We presented MorphVAE, an efficient and unsupervised generative model of neural morphologies based on 3Dwalks. On multiple diverse datasets MorphVAE was able to generate new morphologies from reference neurons with a controlled degree of variation that matched key characteristics of their biological counterparts without explicitly incorporating biological constraints. Additionally, MorphVAE yielded a representation for neuron morphologies that integrated label information in a semi-supervised fashion and that was at least as good as commonly used density maps in distinguishing different cell types.

### 4.1. Related Work

Learning representations from raw morphological data has only been explored in one further study so far. Zhang et al. (2021) recently trained LSTMs on morphological reconstruction files coupled with convolutional networks on density maps to successfully classify several types of rat neurons. Their model does not allow for the generation of new data, however. Here, different approaches have been suggested in the past, which we will briefly review below.

Sampling based methods, for example, start with a simple morphological shape and make small changes iteratively using e.g. Markov-Chain Monte-Carlo (MCMC) methods (Serene, 2013; Farhoodi & Kording, 2018). These methods are computationally extremely expensive, especially if the morphologies are large. Similarly, mechanistic growth models actively grow neurites from the soma outwards using biologically plausible operations which are triggered according to preset rules or by ‘environmental cues’ (van Pelt & Schierwagen, 2004; Memelli et al., 2013; Torben-Nielsen & De Schutter, 2014; Kassraian-Fard et al., 2020). These methods can be reasonably efficient and allow inference about biological processes, but choosing their parameters is typically not straightforward if one wants to achieve convincing neuron shapes.

Hierarchical models grow neuron segments iteratively by sampling from estimated morphometric priors that determine e.g. segment length, radius or branch angle. This process is repeated until the entire neuron matches the original neuron or neuron class with respect to a set of selected statistics. This approach has been successfully employed to grow cortical pyramidal neurons (Eberhard et al., 2006), and motor neuron and Purkinje cell dendrites (Palombo et al., 2019) but, again, it needs careful selection over the morphometric priors. Similarly, Cuntz et al. (2011) sample 2D and 3D point clouds from an averaged density map and subsequently connect the sampled points via a modified minimal spanning tree algorithm that incorporates the optimization of cytoplasmic volume, space and conduction time. A similar approach that is based on synaptic target points is taken by Bingham et al. (2020).

Thus, MorphVAE provides an efficient alternative for generating neuron morphologies that can operate on a large set of possibly diverse neuron morphologies, while learning a generative model for all of these jointly. Once trained, we can use MorphVAE to sample an unlimited number of morphologies from a reference neuron without much additional computational cost. Moreover, MorphVAE incorporates cell type diversity and representation learning directly and provides an embedding of neuron morphologies, which can be used for exploratory analysis and visualization.

### 4.2. Limitations

Although the neurons sampled from the MorphVAE model looked realistic even to experts, our model consistently underestimated the average branch angles for all neurons. As this is introduced by the post-hoc clustering of the sampled walks, future work will need to explore other aggregation schemes, and conditional sampling of the walks in 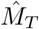. Furthermore, the learned neuron representations cannot be uniquely inverted back into their respective walk matrix which makes meaningful interpolation between different morphologies difficult. A possible remedy could be the interpolation between matched walks in the encoded latent space Z, similar to what has been proposed by Batabyal et al. (2020).

### 4.3. Future work

The MorphVAE model allows to generate diverse sets of neurons with a controlled amount of variation which will be valuable for large-scale simulations and network analysis. It may further generate ground truth data to assess and improve reconstruction algorithms for light- and electron microscopy data (Peng et al., 2010; Helmstaedter et al., 2013; Bria et al., 2016). Finally, through the incorporated representation learning, MoRpHVAE will facilitate further research into the morphological diversity of cell types in the brain, which form the building blocks of the neural circuits underlying perception, decision making, memory and motor action. As MorphVAE learns a generative distribution of plausible and possible walks through neural reconstructions it might also help to detect morphological anomalies in development or neurological diseases.

## Acknowledgements

This work was supported by the German Federal Ministry of Education and Research (BMBF) through the Bernstein Award (01GQ1601) and the Tübingen AI Center (FKZ: 01IS18039A), and the German Research Foundation (DFG) through a Heisenberg professorship (BE5601/4-1) and the Excellence Strategy EXC 2064/1 (project number 390727645).

## Footnotes

1 https://download.brainimagelibrary.org/3a/88/3a88a7687ab66069/

