## Appendix for "MorphVAE: Generating Neural Morphologies from 3D-Walks using a Variational Autoencoder with Spherical Latent Space"

### A. Data

#### A.1. Data generation

We generated a set of toy neurons where each neuron iteratively grows an arbors into one of the four cardinal directions (up, down, left, right). Here, we will describe the procedure for the right arbor, other arbors are generated equivalently. For each neuron we started with one active node at the soma. For each active node $v_j$ we sampled the number of outgoing branch segments $bo_j \sim \max(1, Poisson(\lambda_i))$ from a quasi Poisson distribution. For each outgoing segment we sampled its direction as an angle $\gamma_j$ to the Cartesian basis vector $b = (1, 0, 0)$ with $\gamma_j \sim \mathcal{U}(\alpha_i, \beta_i)$ being drawn from a uniform distribution over the minimal and maximal path angle $\alpha_i$ and $\beta_i$ respectively. The sampled directed segments were then appended to the coordinate of the active node with a segment length of 1, their end points become new active nodes and the former active node $v_j$ becomes inactive. This process was repeated for all active nodes until the arbor contained a specified number of segments.

##### A.1.1. Artificial dataset

Following this procedure we generated $N = 1200$ artificial neurons with $|V| = 200$ nodes that belonged to one of three different populations $P_i$ of equal size ($N_{P_i} = 400$). Each population grew into the horizontal direction (having a right and a left arbor, with 99 segments each) and had its unique set of parameters $\{\lambda_{P_1} = 1, \alpha_{P_1} = -50, \beta_{P_1} = 50\}$, $\{\lambda_{P_2} = 2, \alpha_{P_2} = -50, \beta_{P_2} = 50\}$, and $\{\lambda_{P_3} = 3, \alpha_{P_3} = -100, \beta_{P_3} = 80\}$.

##### A.1.2. Artificial dataset for pre-training

We also generated $N = 1500$ artificial neurons with $|V| = 400$ nodes that belonged to one of five different populations $P_i$ of equal size ($N_{P_i} = 300$). Each populations had its unique set of parameters $\{dir_{P1} = [left, right], \lambda_{P_1} = 1, \alpha_{P_1} = -50, \beta_{P_1} = 50\}$, $\{dir_{P2} = [left, right], \lambda_{P_2} = 2, \alpha_{P_2} = -80, \beta_{P_2} = 80\}$, $\{dir_{P3} = [left, right, up, down], \lambda_{P_3} = 2, \alpha_{P_3} = -80, \beta_{P_3} = 80\}$, $\{dir_{P4} = [up, down], \lambda_{P_4} = 1, \alpha_{P_4} = -50, \beta_{P_4} = 50\}$, and $\{dir_{P5} = [up, down], \lambda_{P_5} = 2, \alpha_{P_5} = -80, \beta_{P_5} = 80\}$. This dataset was used for pre-training the models as it better matched the walk lengths and the walk directions that were encountered in the real datasets.

| Cell type | Names from Bae et al. (2018) | | Count |
|---|---|---|---|
| clus1 | 1wt | sOFF$\alpha$ | 121 |
| clus2 | 2an | F-mini OFF | 29 |
| clus3 | 27 | – | 11 |
| clus4 | 37c | ON-OFF | 17 |
| clus5 | 4i/4on | mini tOFF$\alpha$ | 44 |
| clus6 | 4ow | tOFF$\alpha$ | 91 |
| clus7 | 5to | – | 20 |
| clus8 | 5si | HD1/HD2 | 37 |
| clus9 | 63 | F-mini ON | 19 |
| clus10 | 6sn | – | 36 |
| clus11 | 6sw | tON$\alpha$ | 21 |
| clus12 | 6t | F-midi ON | 30 |
| clus13 | 8w | sON$\alpha$ | 44 |
| clus14 | 8n/9n | – | 30 |
| -1 | 1ni | OFF Step | 4 |
| -2 | 2o | G5bc | 1 |
| -3 | 25 | – | 4 |
| -4 | 28 | – | 5 |
| -5 | 3i | – | 3 |
| -6 | 3o | – | 1 |
| -7 | 51 | W3B | 4 |
| -8 | 72 | – | 1 |
| -9 | 73 | OND | 4 |
| -10 | 81i | – | 2 |
| -11 | 82n | – | 5 |
| -12 | 82wi | vOS | 1 |
| -13 | 85 | – | 3 |
| -14 | 9w | M2 | 1 |
| -15 | 91 | – | 3 |
| -16 | 915 | – | 3 |
| -17 | Fbistrat | F-bistratified | 4 |

*Table 1.* Cluster names and their cell numbers for the retinal ganglion cell dendrites. All clusters with negative cluster number were used for training the autoencoder but not for classification as their numbers were too small.

#### A.2. Pyramidal neurons from motor cortex

We downloaded 275 dendritic reconstructions of excitatory neurons[1] that had been recorded in a large scale multi-modal study describing cell types in adult mouse M1 (Scala et al., 2020). The morphological cell type (m-type) of these excitatory neurons was manually assigned to one of tufted, untufted or other based on visual inspection of the apical dendrites. 'Tufted' neurons showed a small or big apical tuft with at least three tips reaching towards the direction of layer 1. 'Untufted' neurons showed a single apical dendrite with no tuft. All other neurons like star-shaped stellate cells, or neurons with inverted or

[1] https://download.brainimagelibrary.org/3a/88/3a88a7687ab66069/excitatory/

horizontal dendrites were labelled as ‘others’. This yielded a total of $n_t = 135$ tufted, $n_u = 107$ untufted, and $n_o = 33$ other neurons.

##### A.3. Inhibitory neurons from motor cortex

We downloaded 372 axonal reconstructions of inhibitory neurons[2] that had been recorded in a large scale multi-modal study describing cell types in adult mouse M1 (Scala et al., 2020). We used the assigned RNA family labels (Sst, Pvalb, Vip, Sncg, and Lamp5) as cell type labels but grouped Sncg to Vip as it contained only 6 cells. This yielded a total of $n_{Sst} = 108$, $n_{Pvalb} = 145$, $n_{Vip} = 69$, and $n_{Lamp5} = 47$ neurons. 3 neurons had a mismatch in RNA label. We used their reconstructions for training the autoencoder but ignored their labels for classification.

##### A.4. Retinal ganglion cell dendrites

We downloaded 599 reconstructions of retinal ganglion cell dendrites from neuromorpho (Ascoli et al., 2007) that were originally collected by Reinhard et al. (2019) and reconstructed using the TREES Toolbox (Cuntz et al., 2011) using a balancing factor of .4. We used the cell type labels assigned by the authors which yielded 550 cells grouping into 14 classes with at least 11 cells per class. The remaining 49 cells mapped to clusters with lower cell numbers (see Table 1). We used these reconstructions for training the autoencoder but ignored their labels for classification.

##### A.5. Generating image stacks

We compared our generative model with the TREES Toolbox (Cuntz et al., 2011). To sample new morphologies, the Toolbox requires 3D image stacks, however. For this, we sampled points along the neurites of each neuron at a distance of 1 micron and binned the resulting point cloud into 128 bins in each direction. Hereby, each neuron was normalized locally to increase the signal in each image stack. The individual voxel sizes were passed to the Toolbox to allow for proper scaling during the reconstruction. To introduce variation, the image stacks were smoothed using a Gaussian filter with $\sigma = 2$ as implemented in `scipy.ndimage.gaussian_filter`. The image stacks were then uploaded to MATLAB Online and a custom MATLAB script was run that used the functionality of the TREES Toolbox to sample one new morphology per stack.

#### B. Representations

##### B.1. Density maps

We generated several density maps to compare our learned representation to. Table 2 shows the parameters used for density map generation. The normalization ranges were determined as the rounded down $5th$-percentile and the rounded up $95th$-percentile of all reconstructions' point coordinates within each dataset.

| Dataset | Projection | $\sigma$ | $n_{PC} > 95\%$ var | Ranges |
|---|---|---|---|---|
| Toy | XY | .5 | 105 | $x = [-5, 5]$ |
| | | 1 | 26 | $y = [-5, 5]$ |
| | | 2 | 10 | $z = [-5, 5]$ |
| M1 EXC | XZ | .5 | 64 | $x = [-3, 2]$ |
| | | 1 | 22 | $y = [-4, 2]$ |
| | | 2 | 7 | $z = [-3, 9]$ |
| M1 INH | XZ | .5 | 28 | $x = [-4, 4]$ |
| | | 1 | 9 | $y = [-3, 2]$ |
| | | 2 | 4 | $z = [-6, 6]$ |
| RGC | Z | .5 | 6 | $x = [-6, 4]$ |
| | | 1 | 4 | $y = [-4, 5]$ |
| | | 2 | 3 | $z = [-1, 1]$ |

*Table 2.* Summary of the parameters used for density map generation. It shows the projection axes, the smoothing kernels $\sigma$, the number of principle components that have been kept and the normalization ranges for each direction.

##### B.2. Morphometric features

We computed morphometric statistics using the default functionality of the `MorphoPy` toolbox (Laturnus et al., 2020). This yielded a total of 28 single valued morphometric statistics for each dataset. Since we did not model the morphologies' thickness, we excluded the four morphometric statistics that related to that feature in each dataset, namely, average thickness, maximal thickness, total surface, and total volume. Additionally, we excluded all statistics that were zero for most of the cells, namely, log minimal tortuosity and log median tortuosity. The remaining 22 features are shown for the artificial data (see Fig. B.2) and the real datasets (see Fig. B.3, Fig. B.4, and Fig. B.5), their respective samples for different

[2] https://download.brainimagelibrary.org/3a/88/3a88a7687ab66069/inhibitory/

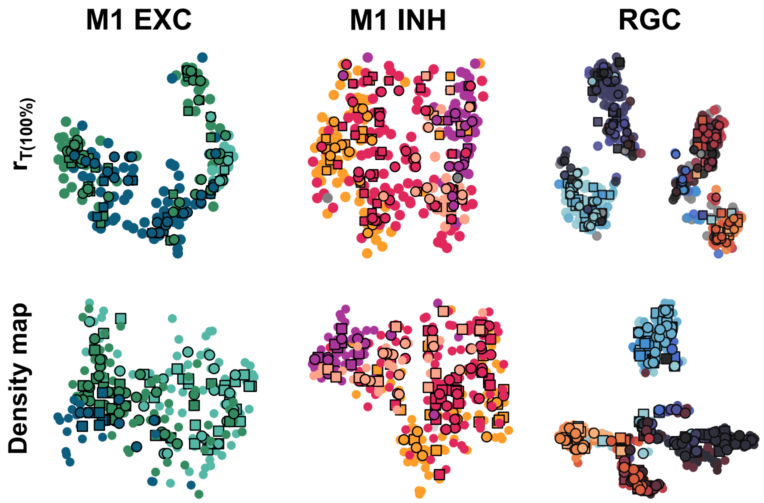


*Figure B.1.* T-SNE embedding of the learned neural representation $r_T$ and the best performing density map. For labels see Fig. B.3 – Fig. B.5.

$\kappa$, and the comparison with the TREES Toolbox (Cuntz et al., 2011). Note, for the inhibitory cell data we also excluded the soma exit angle feature as they only exhibit one axonal neurite.

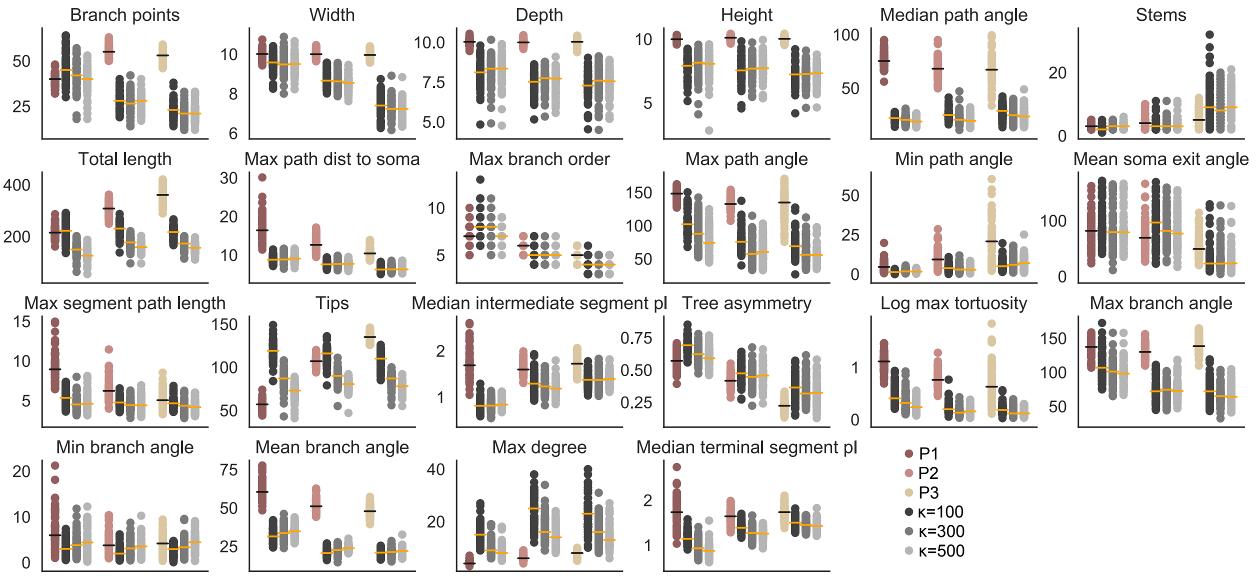


*Figure B.2.* Distributions of all computed morphometric statistics for the test neurons in each population (colored) of the artificial dataset and the sampled neurons using different values of $\kappa$ during sampling in the latent space (grey). Lines indicate the medians.

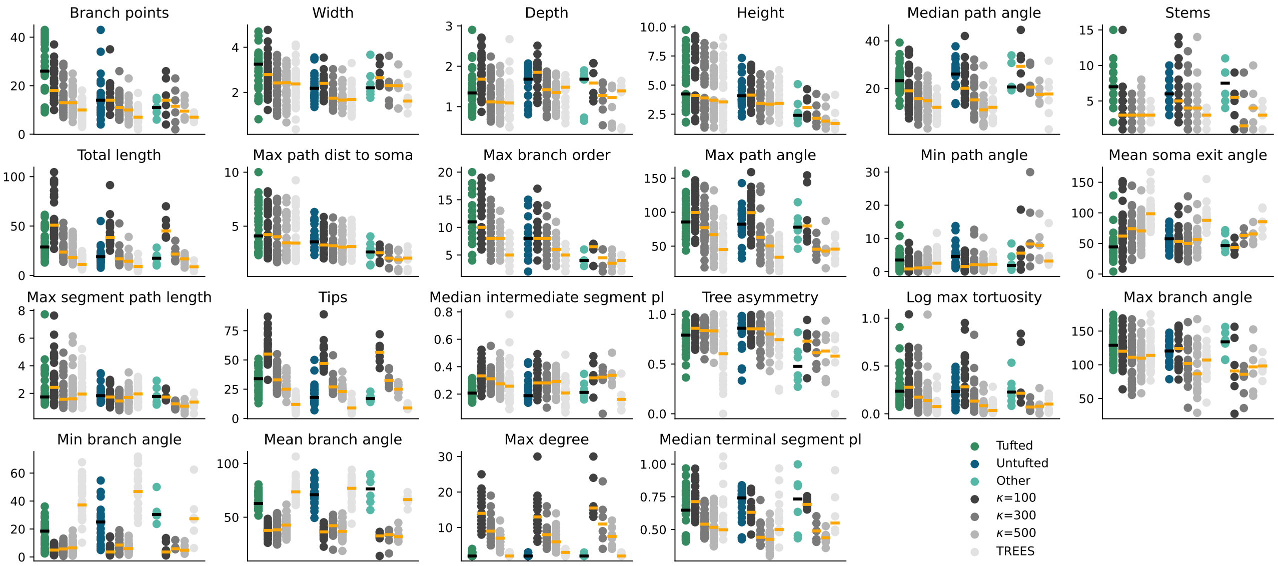


*Figure B.3.* Distributions of all computed morphometric statistics for the pyramidal cell dataset. The data is shown for the test neurons in each population (colored), the sampled neurons using MORPHVAE with different values of $\kappa$ during sampling in the latent space (grey), and sampled neurons using the TREES Toolbox (light grey). Lines indicate the medians.

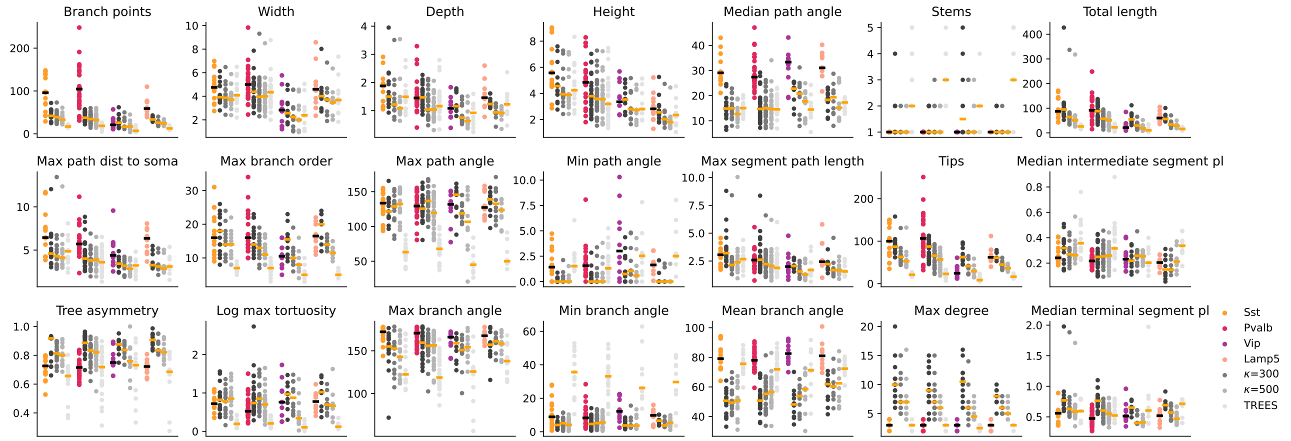


*Figure B.4.* Distributions of all computed morphometric statistics for the inhibitory cell dataset. The data is shown for the test neurons in each population (colored), the sampled neurons using MORPHVAE with different values of $\kappa$ during sampling in the latent space (greys), and sampled neurons using the TREES Toolbox (light grey). Lines indicate the medians.

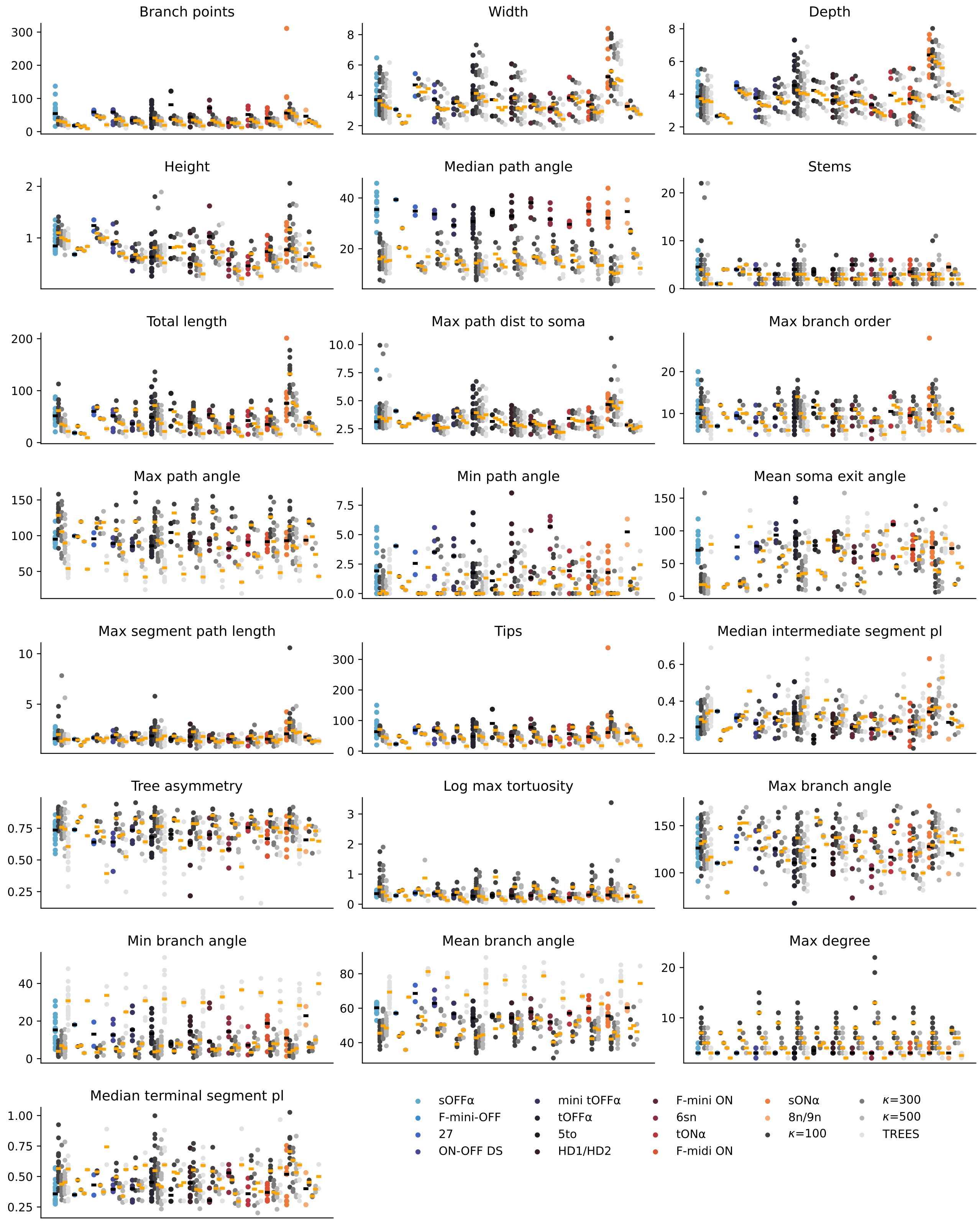


*Figure B.5.* Distributions of all computed morphometric statistics for the retinal ganglion cell dataset. The data is shown for the test neurons in each population (colored), the sampled neurons using MORPHVAE with different values of $\kappa$ during sampling in the latent space (greys), and sampled neurons using the TREES Toolbox (light grey). Lines indicate the medians.

### C. Model

#### C.1. Model architecture

The MorphVAE model is inspired by previous work in natural language processing (Sutskever et al., 2014; Xu & Durrett, 2018; Davidson et al., 2018) but has been adjusted to predict continuous variables instead of discrete word tokens. Fig. C.1 shows a more detailed account of the model architecture.

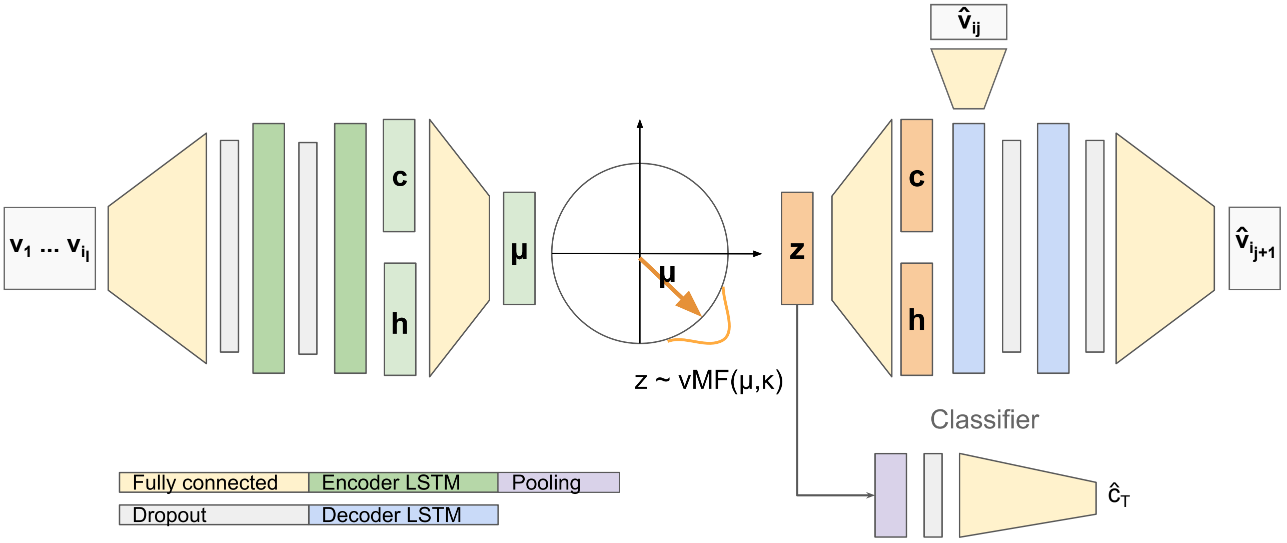


*Figure C.1.* A detailed depiction of all model components. The encoder and the decoder are both two-layered unidirectional LSTMs. Each walk $w = (v_1, \cdots, v_{i_l})$ encodes a the mean $\mu$ of a von-Mises Fisher distribution which is used to sample a latent representation $z$. The latent representation $z$ is then used to initialize the hidden and the cell state, $h$ and $c$, of the decoder, that will predict the next node coordinate $\hat{v}_{i_j}$ from its internal states and the previous prediction $\hat{v}_{i_j}$. Additionally, a pooling layer aggregates all latent representations $z_i$ that describe the walks within one neuron $T$ and predicts the neuron's cell type label $\hat{c}_T$.

#### C.2. Model selection

To find the optimal values for the network dimensions and pooling operations we performed a grid search over $m = [16, 32]$, $k = [8, 16, 32]$, dropout $= [.1, .3, .5]$, $\kappa = [100, 500]$ and $Pool = [\text{max-pooling, average-pooling}]$. We took the model with the best average validation performance over three Glorot initializations (Glorot & Bengio, 2010). The best model used a hidden and a latent dimension of $m = k = 32$, dropout of .1, a variance of $\kappa = 500$ and max-pooling.

#### C.3. Runtime analysis

| | TREES Toolbox | | | MorphVAE | | |
|---|---|---|---|---|---|---|
| Dataset | Loading image stack | Sampling points | MST | Encoding walk matrices | Sampling new walks | Clustering |
| M1 EXC | 16.59s ± 1.73 | 0.38s ± 0.08 | 0.18s ± 0.04 | 6.04s ± 0.19 (2.38s ± 0.01) | 0.65s ± 0.03 (0.13s ± 0.0) | 0.02s ± 0 |
| M1 INH | 16.19s ± 0.75 | 0.39s ± 0.05 | 0.22s ± 0.04 | 6.34s ± 0.89 (2.47s ± 0.01) | 0.72s ± 0.03 (0.12s ± 0.0) | 0.03s ± 0.0 |
| RGC | 16.61s ± 0.95 | 0.43s ± 0.4 | 0.28s ± 0.54 | 7.94s ± 0.56 (3.96s ± 0.0) | 0.71s ± 0.06 (0.13s ± 0.0) | 0.05s ± 0.0 |

*Table 3.* Wall clock runtime in seconds for different operations during morphology generation for the TREES Toolbox and MorphVAE. Values in brackets denote the runtime on the GPU.

We compared the runtime during morphology sampling for the MorphVAE model and the TREES Toolbox (Cuntz et al., 2011) in MATLAB Online (Table 3). For comparability we restricted our system to the same amount of CPUs and disabled GPU processing. A more detailed description of each system is given in Table 4. The runtime during sampling and tree construction was similar for both models (Table 3) but the TREES Toolbox needs to load a new image stack for each neuron which slows the run time down considerably. In comparison, MorphVAE encodes the reference walk matrices once for each batch and its implementation allows for GPU processing which leads to substantial runtime improvements.

| | MATLAB Online | Our setup |
|---|---|---|
| Architecture | x86_64 | x86_64 |
| CPU op-mode(s) | 32-bit, 64-bit | |
| CPU(s) | 16 | 16 |
| Thread(s) per core | 2 | 2 |
| Core(s) per socket | 8 | 10 |
| CPU family | 6 | 6 |
| Model | 85 | 79 |
| Stepping | 7 | 1 |
| CPU MHz | 3100.310 | 1200.036 |
| BogoMIPS | 4999.99 | 4399.83 |
| Virtualization type | full | VT-x |
| L1d cache | 32K | 32K |
| L1i cache | 32K | 32K |
| L2 cache | 1024K | 256K |
| L3 cache | 36608K | 25600K |

*Table 4.* Systems description of MATLAB Online and our setup when executing *lscpu*.
